# Effect of a *Monascus* sp. red yeast rice extract on germination of bacterial spores

**DOI:** 10.1101/2021.03.26.437149

**Authors:** Husakova Marketa, Plechata Michaela, Branska Barbora, Patakova Petra

## Abstract

The pink-red colour of traditional sausages (cured meat) is the result of nitrite addition and the formation of nitrosomyoglobin. However, the pleasant colour of processed meat products is a side effect of nitrite addition while the main anticipated goal is to suppress the germination of clostridial spores. The fungus *Monascus* is known as a producer of oligoketide pigments, which are used in Asian countries, especially in China, for colouring foods, including meat products. Although, different biological activities of *Monascus* pigments have been tested and confirmed in many studies, their effect on germination of bacterial spores has never been investigated. This study is focused on testing the activity of red yeast rice (RYR) extract, containing monascin, rubropunctatin, rubropunctamine complexes and monascuspiloin as the main pigments, on germination of *Clostridium* and *Bacillus* spores. It was found that addition of nitrite alone, at the permitted concentration, had no effect on spore germination. However, the combined effects of nitrite with NaCl, tested after addition of pickling salt, was efficient in inhibiting the germination of *C. beijerinckii* spores but had no effect on *B. subtilis* spores. In contrast, total suppression of *C. beijerinckii* spore germination was reached after addition of RYR extract to the medium at a concentration of 2 % v/v. For *B. subtilis*, total inhibition of spore germination was observed only after addition of 4 % v/v RYR extract to the medium containing 1.3% w/w NaCl.

## 1. Introduction

Red yeast rice (RYR) is rice fermented by the fungus *Monascus*, which is prepared for different applications using different *Monascus* species (for a recent review see Zhu et al., 2019). RYR has various synonyms such as hong-qu, beni-koji or ang-kak in languages of Asian countries, where the product is popular. In Europe, RYR is only permitted as a food supplement on the condition that it is prepared using *Monascus purpureus* and the preparation (food supplement) should contain 10 mg of monacolin K, administered daily, in order to guarantee the effect as described in the health claim “Monacolin K from red yeast rice contributes to the maintenance of normal blood cholesterol levels” (Commission Regulation (EU) No 432/2012). As a dose of 10 mg of monacolin K corresponds to the lowest therapeutically effective dose of statins in prescription drugs, the required amount of monacolin K in RYR food supplements was reconsidered by the EFSA Panel on Food Additives and Nutrient Sources added to Food (ANS) in 2018 (EFSA Panel, 2018) but with no clear conclusion. Nevertheless, the main concern associated with the use of RYR is its potential contamination with citrinin, a mycotoxin whose toxicological effects on people have not been fully elucidated (de Oliveira Filho et al., 2017). By the Commission Regulation (EU) 2019/1901, the maximum tolerated citrinin concentration in RYR was set to 100 µg/kg.

The fungus *Monascus* is especially known for its production of red pigments, which are used in certain Asian countries, such as China, Japan or Philippines, for food colouring. As the colour red is associated with many different fruits, vegetables and meat products, *Monascus* pigments are mostly used for colouring cakes or other sweet products, fruit yoghurts or other fermented milk products and processed meat. The suitability of *Monascus* pigments for colouring meat products, particularly with regard to colour, texture, smell and other sensory parameters of the products, has already been proven in the scientific literature (Leistner et al., 1991; Fabre et al., 1993; Yu et al., 2015; Seong et al., 2017). In addition, inhibitory effects of *Monascus* pigments or *Monascus* extracts on vegetative bacterial cells e.g. *Staphylococcus aureus, Escherichia coli, Bacillus subtilis* ((Kim et al., 2006; Vendruscolo et al., 2014; Zhao et al., 2016) have been demonstrated. In addition, the safety of *Monascus* pigments for human consumption has been confirmed in several studies (Bianchi, 2005; Yu et al., 2008; Mohan Kumari et al., 2009).

The aim of the study was to test whether an ethanol extract of red yeast rice, having a red colour and containing a mixture of *Monascus* pigments but without citrinin and monacolin K, might suppress germination of bacterial spores. In traditional food processing, nitrite salts have been added to meat products in order to achieve total inhibition of germination of *Clostridium botulinum* spores. Nitrite salts are also responsible for the pleasing red colour of processed meat products, caused by the formation of nitrosomyoglobin. In the work described for the first time here, *Clostridium beijerinckii* and *Bacillus subtilis* spores were used as models of anaerobic and aerobic bacterial spore formers, respectively.

## 2. Material and Methods

### 2.1 RYR preparation

*Monascus* sp. DBM 4361, isolated from a nonsterile dried red fermented rice sample, was maintained on Potato-Dextrose agar (VWR Chemicals) slants at 4°C. The strain was deposited at the Department of Biochemistry and Microbiology (DBM), University of Chemistry and Technology Prague.

An amount of rice (Giana, Thailand) (150 g) was washed with hot water, then boiled for 1 minute. The rice was evenly divided into three autoclavable plastic bags, which were closed with a metal ring and a cotton plug. The bags were placed in a beaker sealed with aluminium foil and sterilized at 121 °C for 20 minutes. Sterilization was repeated after 24 hours to eliminate contamination by spore-forming bacteria. Spores from the *Monascus* culture (mixture of ascospores and conidia, because the strain formed both asexual and sexual spores, see Fig. 1) were transferred to sterile water using a sterile loop. The sterile rice in the bags was inoculated with 5 mL of the spore suspension. Cultivation of the fungus on rice was performed for 10 days at 30 °C. The rice was mixed by hand daily.

**Fig. 1.**
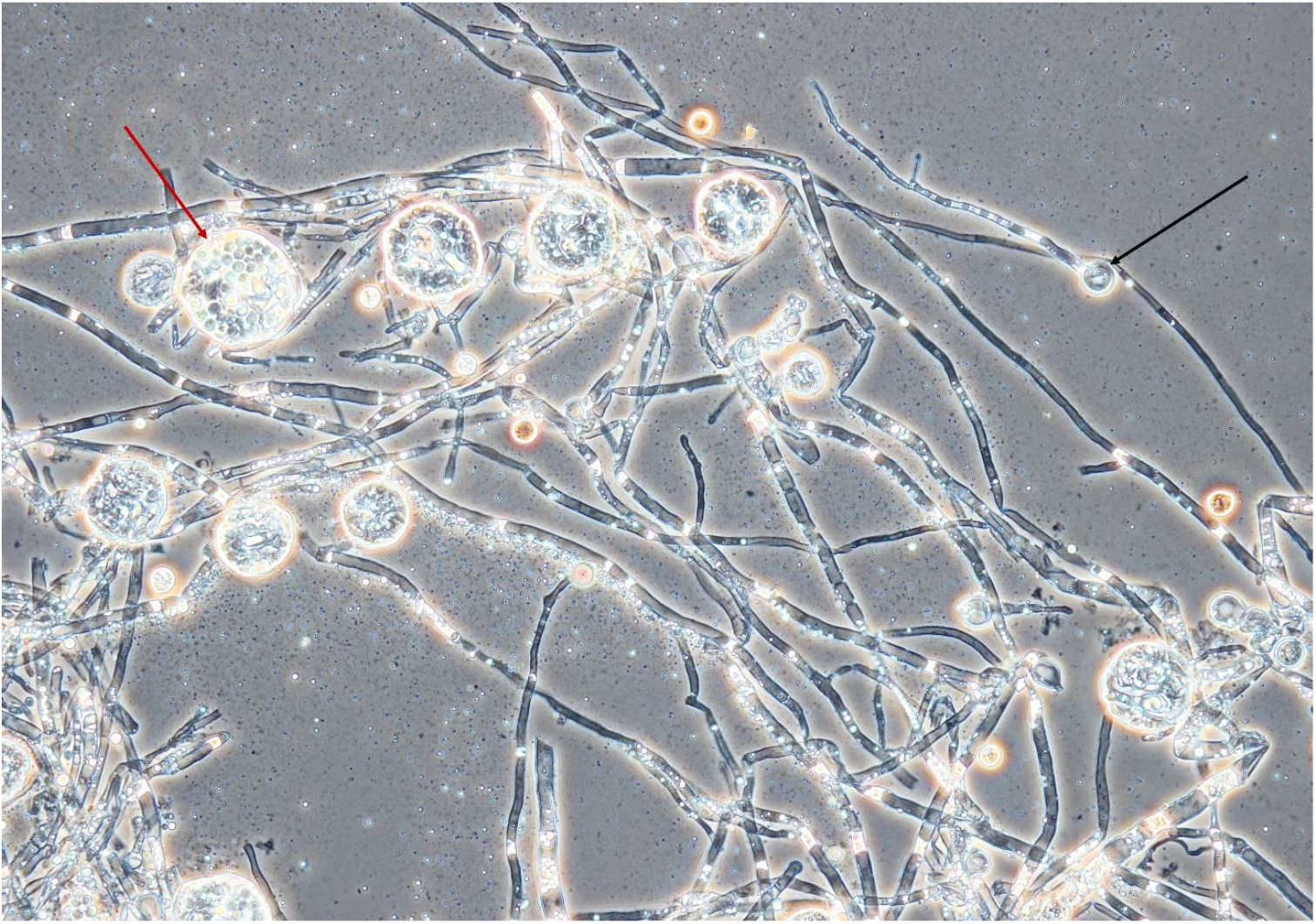
Mycelium, conidia and cleistothecia of *Monascus* sp. DBM 4361. An example of a conidium is marked with a black arrow, the red arrow shows a cleistothecium with ascospores. The specimen was prepared from the fungus grown on a PDA agar slant for 5 days at 30 °C; magnification was 400x.

### 2.2 Extraction of pigments, pH estimation

The RYR (Fig.2) (5 g) was extracted with 25 mL of 70% ethanol and distilled water in 250mL Erlenmeyer flasks for 1 h, at 30°C, with shaking (laboratory shaker Infors, 100 rpm). The mixture was then filtered through Whatman 1 filter paper. Pooled ethanol extracts from 3 flasks were concentrated using a rotary vacuum evaporator (Boeck) (max. temperature 55 °C), so that all ethanol was evaporated. The remaining water extract contained insoluble pigmented particles, which were collected by filtration and dissolved in 96% ethanol. The resulting ethanol extract was used for all microbiological assays and was analyzed by HPLC. The pH of pooled water extracts from 3 flasks was measured and shown to be 4.9.

**Fig. 2.**
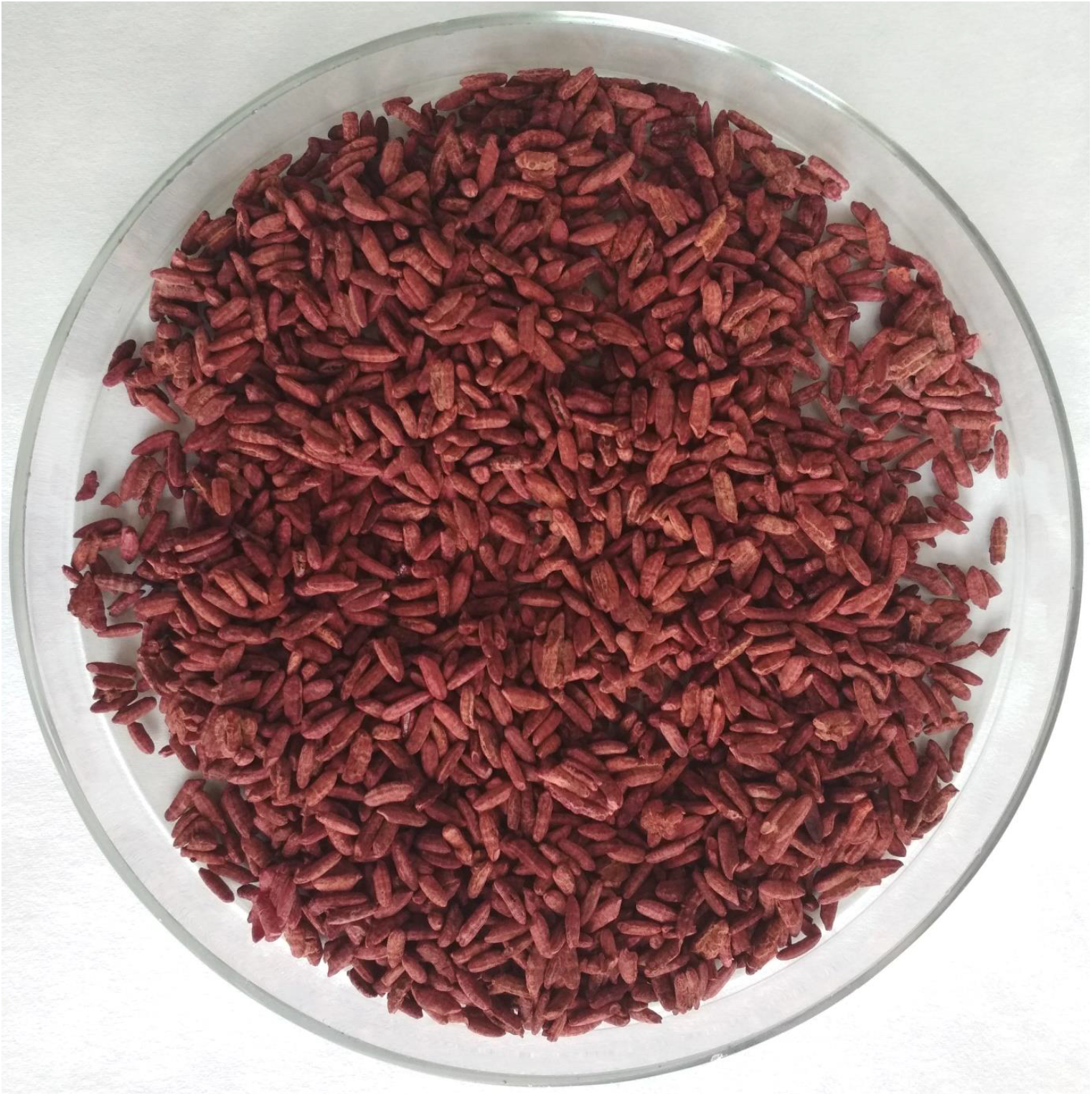
Red yeast rice fermented with *Monascus* sp. DBM 4361 for 10 days at 30 °C.

### 2.3 RYR extract analysis

#### 2.3.1 Spectrophotometric analysis of the RYR extract

The RYR extract was diluted 200-fold with 96% ethanol to adjust the absorbance to 0.1-1.0 at 330 – 600 nm. The absorbance of the sample was detected using a spectrophotometer (Varian Cary 50 Bio). The results were represented as an absorbance spectrum or as an absorbance value at a selected wavelength, the dilution factor being taken into consideration. As a blank, 96% ethanol was used.

#### 2.3.2 UHPLC analysis of pigments, citrinin and monacolin K

UHPLC (Agilent Technologies 1260 Infinity II) was used to determine *Monascus* pigments, citrinin and monacolin K. The following conditions were used: Kinetex Polar C18, 100A, 150 x 4.6 mm column; the mobile phase: 0.025% H_3_PO_4_ in water:acetonitrile at a ratio of 60:40; isocratic elution at a flow rate 1.5 mL/min; injection volume 5 µL. For determination of yellow, orange and red pigments, a photodiode detector set at 390, 470 and 500 nm resp. was used. The presence of monacolin K was detected at 237 nm. For the determination of the mycotoxin citrinin, the fluorescence detector setting was 331 nm for excitation and 500 nm for emission. For analysis, the extract sample was diluted 10-fold with 96% ethanol.

The standards of mycotoxin citrinin (Sigma-Aldrich), yellow pigment monascin (Sigma-Aldrich), and orange pigment rubropuctatin (1717 CheMall Corporation) were used as reference samples. A rubropunctamine laboratory standard was prepared from the rubropunctatin standard by reaction with NH_4_OH (Penta). Unknown yellow, orange and red pigments were identified based on their absorption spectra and quantified as equivalents to their respective standards, i.e. monascin, rubropunctatin and rubropunctamine. For identification of individual compounds in the chromatogram, previous results (Patrovsky et al., 2019) were used.

#### 2.3.3 HPLC-MS analysis

HPLC-HRMS (Accella 600 Thermo Scientific) was used to determine molecular weights of the *Monascus* pigments. The following conditions were used: Luna Omega Polar 1.6 µm, 50 x 2.1 mm Phenomenex column; mobile phase: 0.1% HCOOH in water:methanol; gradient elution (A:B in ratio from 90:10 to 5:95) at a flow rate of 300 µL/min; injection volume 5 µL; ESI-positive mode; LTQ Orbitrap Velos mass analyzer (Thermo Scientific). For the analysis, the extract sample was diluted 200-fold with a solution of 90 % water 10 % methanol.

### 2.4 Spore germination assays

#### 2.4.1. Preparation of spore suspensions

*Clostridium beijerinckii* NCIMB 8052 and *Bacillus subtilis* DBM 3006 were stored in the form of spore suspensions in sterile distilled water at 4 ^°^C. Spores of *C. beijerinckii* NCIMB 8052 were obtained after 48 h incubation of the culture in 250 mL Erlenmeyer flasks containing 100 mL of TYA medium in an anaerobic chamber (Concept 400, Ruskinn Technology, UK) at 37 °C. TYA medium contained in g/L: glucose 40, yeast extract (Merck) 2, tryptone (Sigma Aldrich) 6, potassium dihydrogenphosphate 0.5, ammonium acetate 3, magnesium sulfate heptahydrate, 0.3, ferrous sulfate heptahydrate 0.01; the pH of the medium was adjusted prior to sterilization in the autoclave (20 min, 121 °C, 0.1 MPa) to 6.8. Spores of *B. subtilis* DBM 3006 were obtained after 48 h incubation of the culture in 250 mL Erlenmeyer flasks containing 50 mL of MP broth shaken on a rotary shaker (New Brunswick Scientific) at 300 rpm and 30 °C. MP broth contained in g/L: meat extract (Roth) 3, peptone (Roth) 5; the pH of the medium was adjusted prior to sterilization in the autoclave (20 min, 121 °C) to 7.0. Salts, HCl and NaOH for medium preparation and pH adjustment were purchased from Penta, Czech Republic. After cultivation, spores of both bacterial cultures were harvested by centrifugation (Hettich MIKRO 220R) for 5 min, 5000 rpm, at 4 °C. Spores were washed with sterile water, centrifuged under the same conditions and re-suspended in 20 mL of sterile water. Finally, the spore suspension was pipetted in 1 mL portions into Eppendorf tubes, which were stored at 4°C. All spore handling was performed under aseptic conditions using sterile tools and materials. Spore concentrations were estimated by flow cytometry (Branska et al., 2018) to be 2.10^8^ spores/mL and 8.10^8^ spores/mL for *C. beijerinckii* and *B. subtilis*, respectively. Prior to inoculation of medium for germination assays, spore suspensions were heat shocked (80 °C for 30 s followed by cooling on ice for 2 min) to accelerate germination.

#### 2.4.2. *C. beijerinckii* germination assay

The cultivation tests were performed in 20 mL test tubes containing 9.8 mL of medium inoculated with 0.2 mL heat shocked spore suspension of *C. beijerinckii*. Unmodified TYA medium was used as a positive control. The cultivation tests were performed in triplicate, in the anaerobic chamber, at 37°C for 48h. Different combinations of agents were added to the TYA medium to test their effect on spore germination; see Table I.

**Table I.**
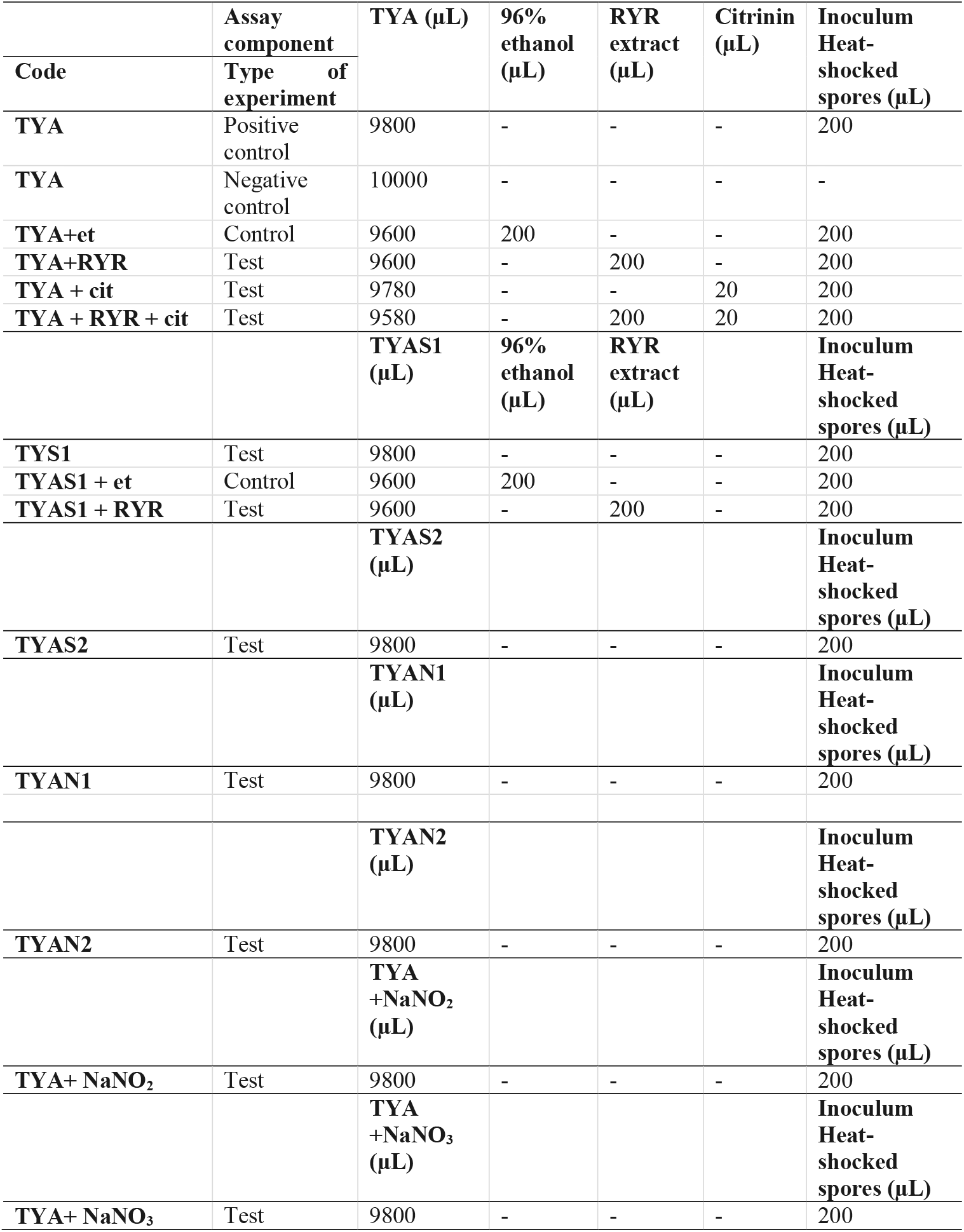
Design of *C. beijerinckii* germination assays in individual test tubes. TYA – unmodified TYA medium with 20 g/L glucose; TYAS1, TYAS2 – TYA medium with the addition of 1.3 % w/w and 2.0 % w/w NaCl, respectively; TYAN1, TYAN2 - TYA medium with the addition of 1.3 % w/w and 2.0 % w/w Praganda butcher’s nitrite pickling mixture (K+S company, Czech Republic); TYA+NaNO_2_ and TYA+NaNO_3_ – TYA media with the addition of 300 mg/L NaNO_2_ or NaNO_3;_ citrinin was added from stock solution, of concentration 100 mg/L in dimethylsulfoxide; et – ethanol, cit – citrinin.

#### 2.4.3. *B. subtilis* germination assay

The cultivation tests were performed on microcultivation plates using the Bioscreen C device (LabSystem) with intermittent shaking (30 seconds every 3 minutes) at 30 °C for 24 h. Each well of the plate was filled with 196 µL of medium and 4 µL of *B. subtilis* heat shocked spore suspension. In each well, optical density was measured at 600 nm, every 30 minutes. Unmodified MP medium was used as a positive control. Each cultivation test was performed in 6 wells. Different combinations of agents were added to the MP medium to test their effect on spore germination, see Table II.

**Table II.**
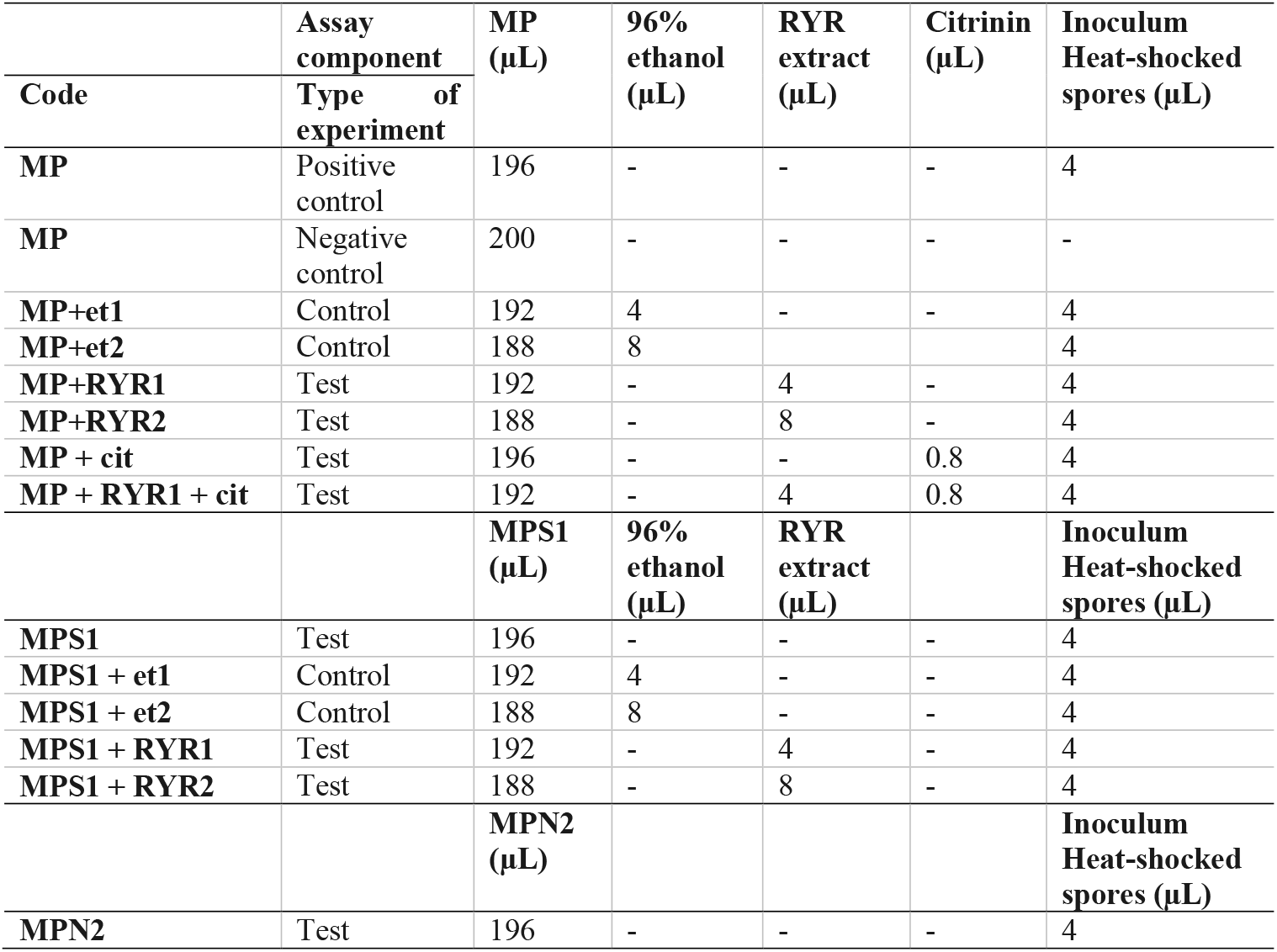
Design of *B. subtilis* germination assays in individual wells of a microcultivation plate. MP – unmodified MP medium; MPS1– MP medium with addition of 1.3 % w/w NaCl; MPN2 – MP medium with addition of 2.0 % w/w Praganda butcher’s nitrite pickling mixture (K+S company, Czech Republic); citrinin was added from a stock solution, of concentration 100 mg/L in dimethylsulfoxide

|  | Assay component | MP (μL) | 96% ethanol (μL) | RYR extract (μL) | Citrinin (μL) | Inoculum Heat-shocked spores (μL) |
| --- | --- | --- | --- | --- | --- | --- |
| Code | Type of experiment |  |  |  |  |  |
| MP | Positive control | 196 | - | - | - | 4 |
| MP | Negative control | 200 | - | - | - | - |
| MP+et1 | Control | 192 | 4 | - | - | 4 |
| MP+et2 | Control | 188 | 8 |  |  | 4 |
| MP+RYR1 | Test | 192 | - | 4 | - | 4 |
| MP+RYR2 | Test | 188 | - | 8 | - | 4 |
| MP + cit | Test | 196 | - | - | 0.8 | 4 |
| MP + RYR1 + cit | Test | 192 | - | 4 | 0.8 | 4 |
|  |  | MPS1 (μL) | 96% ethanol (μL) | RYR extract (μL) |  | Inoculum Heat-shocked spores (μL) |
| MPS1 | Test | 196 | - | - | - | 4 |
| MPS1 + et1 | Control | 192 | 4 | - | - | 4 |
| MPS1 + et2 | Control | 188 | 8 | - | - | 4 |
| MPS1 + RYR1 | Test | 192 | - | 4 | - | 4 |
| MPS1 + RYR2 | Test | 188 | - | 8 | - | 4 |
|  |  | MPN2 (μL) |  |  |  | Inoculum Heat-shocked spores (μL) |
| MPN2 | Test | 196 | - | - | - | 4 |

## 3. Results

### 3.1. *C. beijerinckii* germination

*Clostridium beijerinckii* spores were chosen as a model substituting for *Clostridium botulinum* spores because both species belong to the same Cluster I (*sensu stricto*) of the *Clostridium* genus (Cruz-Morales et al., 2019). Heat-shocked spores of *C. beijerinckii* were inoculated to 2 % by volume to the medium in test tubes and were allowed to germinate under anaerobic conditions. The TYA medium and the inoculation ratio were chosen to guarantee reliable spore germination based on previous experience with the strain (Kolek et al., 2016). The compositions of different media were designed (see Table I) to be able to compare any effect of nitrite salts with the potential effect of the RYR extract, not containing citrinin and monacolin K. Nitrites are only allowed to be added to meat products in a mixture with sodium chloride in the form of NaNO_2_ (E249 food additive) or KNO_2_ (E250 food additive), in an amount not exceeding 150 mg of nitrite per 1 kg of a standard meat product (only in some national specialities produced in different EU countries can the amount of nitrite be higher, up to 300 mg/kg, for certain salamis and bacons and up to 500 mg/kg for herrings and sprouts); for a survey of rules valid in the EU for the addition of nitrite/nitrate to meat products, see Honikel, 2008.

Nitrite, in the form of NaNO_2,_ was added to the culture, either independently or as a component of Praganda nitrite pickling salt. In addition, NaNO_3_ (E251 food additive) was tested because its addition is permitted and applied in cheeses to a maximum concentration of 150 mg/kg, with the aim of suppressing germination of *Clostridium butyricum* spores. While even the addition of NaNO_2_ or NaNO_3_ alone at a concentration of 300 mg/kg had no effect on the germination of *C. beijerinckii* spores, addition of nitrite pickling salt to the recommended concentration for various products i.e. 2 % (w/w) of the pickling salt or 1.3 % (w/w) as recommended for products with low salt content, resulted in total inhibition of spore germination (see Table III, experiment codes TYA+NaNO_2;_ TYA+NaNO_3_; TYAN2 and TYAN1). As *Clostridium beijerinckii* is sensitive to high concentrations of NaCl (Branska et al., 2020), the independent effect of NaCl addition was also tested. While addition of NaCl to 2% w/w suppressed spore germination, 1.3 % w/w did not reliably suppress germination in all cases (see Table III, code TYAS2 and TYAS1).

**Table III.**
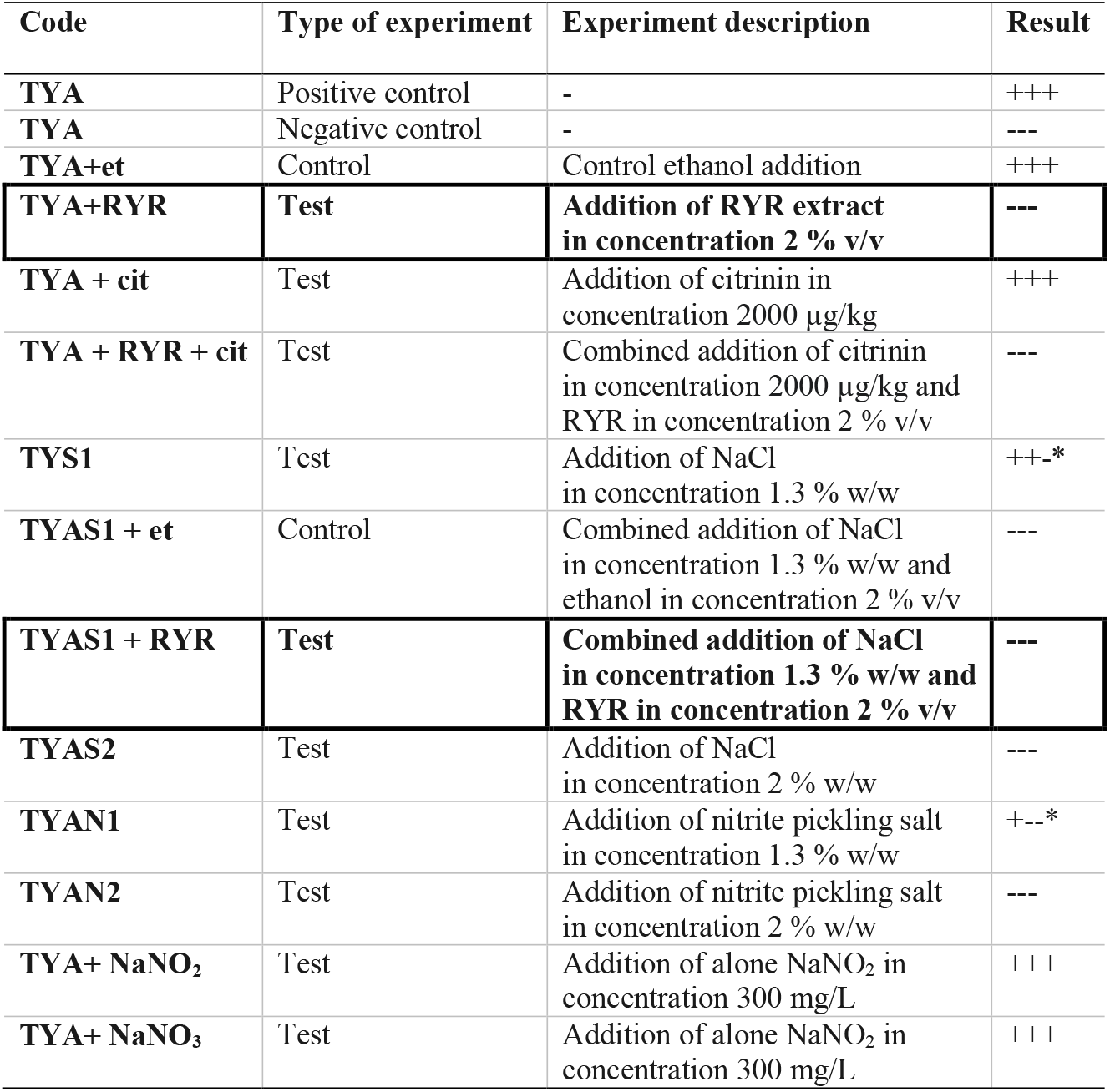
Results of *C*.*beijerinckii* germination assays. The experimental code correlates with the experimental design given in Table I. All experiments were performed in triplicate, spore outgrowth observed in all 3 test tubes is marked as +++, no growth as ---. *In case of dubious results, the cultivation test was repeated but if the result was the same, i.e. ++- or +-- this means unreliable suppression of germination. The most significant results are highlighted.

Addition of 2 % v/v RYR extract was tested with standard TYA medium and with medium containing 1.3 % w/w NaCl; addition of ethanol to the same concentration (2 % v/v) was tested as a control (Table III, codes TYA+RYR, TYA+ Et, TYAS1 +RYR, TYAS1 +Et). While addition of the RYR extract suppressed spore germination, ethanol did not. The RYR extract did not contain citrinin but, because it is known that citrinin has certain antimicrobial properties, the citrinin effect was tested at a concentration of 2000 µg/kg. This concentration of citrinin reflects the amount that was EU-permitted in an RYR food supplement until 2019, after which the limit was reconsidered and adjusted to 100 µg/kg. Citrinin was tested alone or in combination with RYR (Table III, codes TYA+cit, TYA+ RYR + cit).

Typical test tube growth characteristics of *C. beijerinckii* exhibiting high turbidity, foam and development of bubbles of fermentation gas (mixture of CO_2_ and hydrogen) as well as colour of the medium after addition of the RYR extract are shown in Fig.3. Only the ability to grow (indicated as + or -) was tested in the *C. beijerinckii* germination assay. Determination of optical density was not performed in order to not disturb the anaerobic atmosphere. Changed morphology from vegetative cells to spores influences OD values, so comparisons between tests might be misleading. Spores in TYA medium (positive control experiment) started to germinate 16-18 h after inoculation, and in other cases, germination started with different delays.

**Fig. 3.**
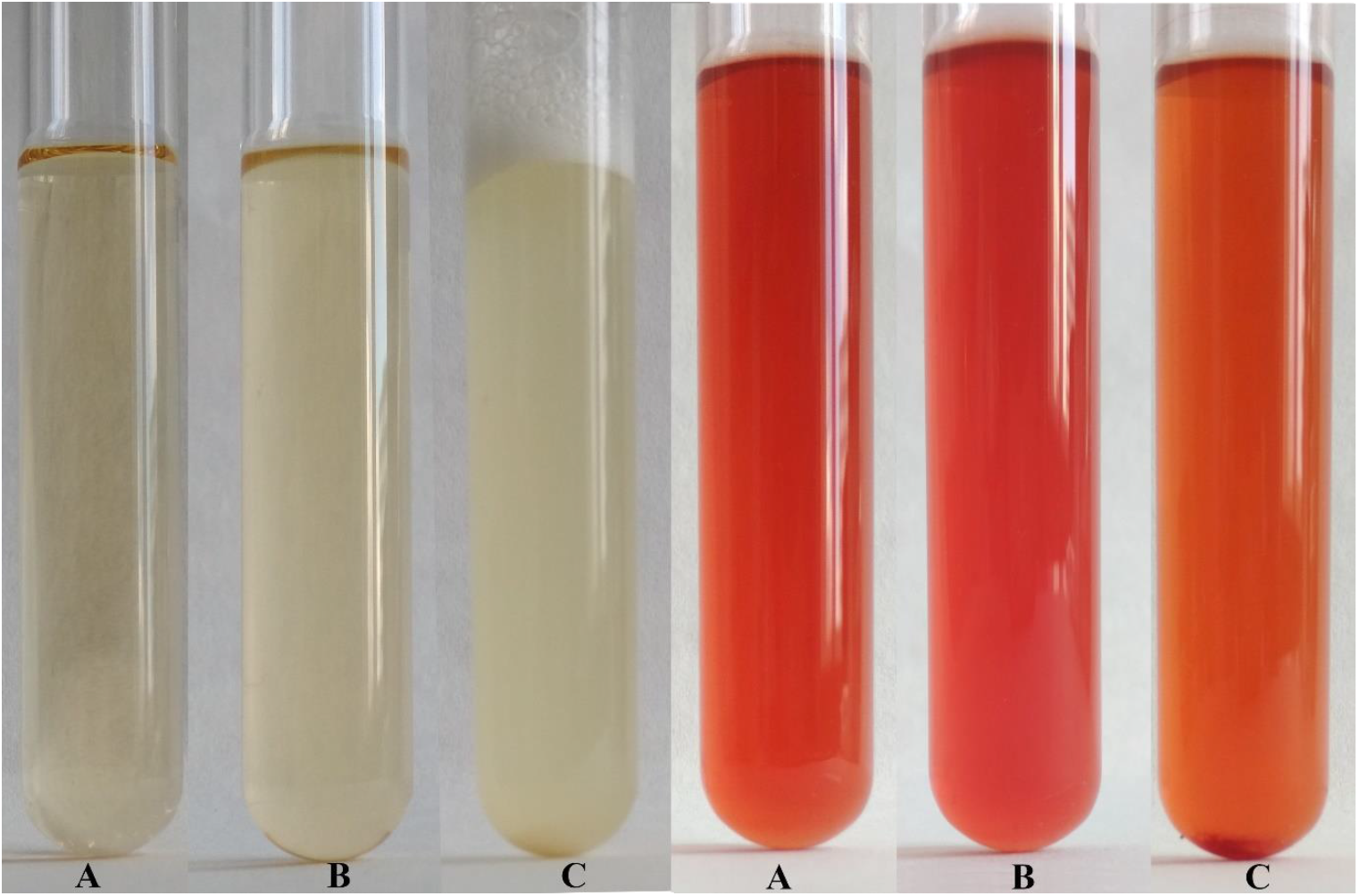
Demonstration of growth in test tubes. Positive control i.e. outgrowth of *C. beijerinckii* spores in TYA medium is shown on the left while suppression of spore germination after addition of RYR extract to 2% v/v is shown on the right. A test tube with medium was photographed prior to inoculation (A), after inoculation (B) and after 48 h cultivation (C)

### 3.2. B.subtilis germination

To follow the growth of spore formers after germination, *Bacillus subtilis* germination assays were performed, even if these spores do not normally occur in meat products (Fig.4). Nevertheless, *B. subtilis* spores are more resistant to adverse environmental effects in comparison with the *C. beijerinckii* spores, therefore the design of experiments had to be different (Table II) in order to inhibit germination. Total suppression of spore germination was achieved only after addition of the RYR extract to 4% v/v in medium containing 1.3 % w/w NaCl (Fig. 4D, MPS1+RYR2). In other cases, growth was always detected, even if, in some cases, germination was delayed by up to 10 h (Fig.4B, MP+ RYR2). Surprisingly, addition of nitrite pickling salt to the recommended 2 % w/w did not inhibit germination of the spores (Fig.4A, MPN2).

**Fig. 4.**
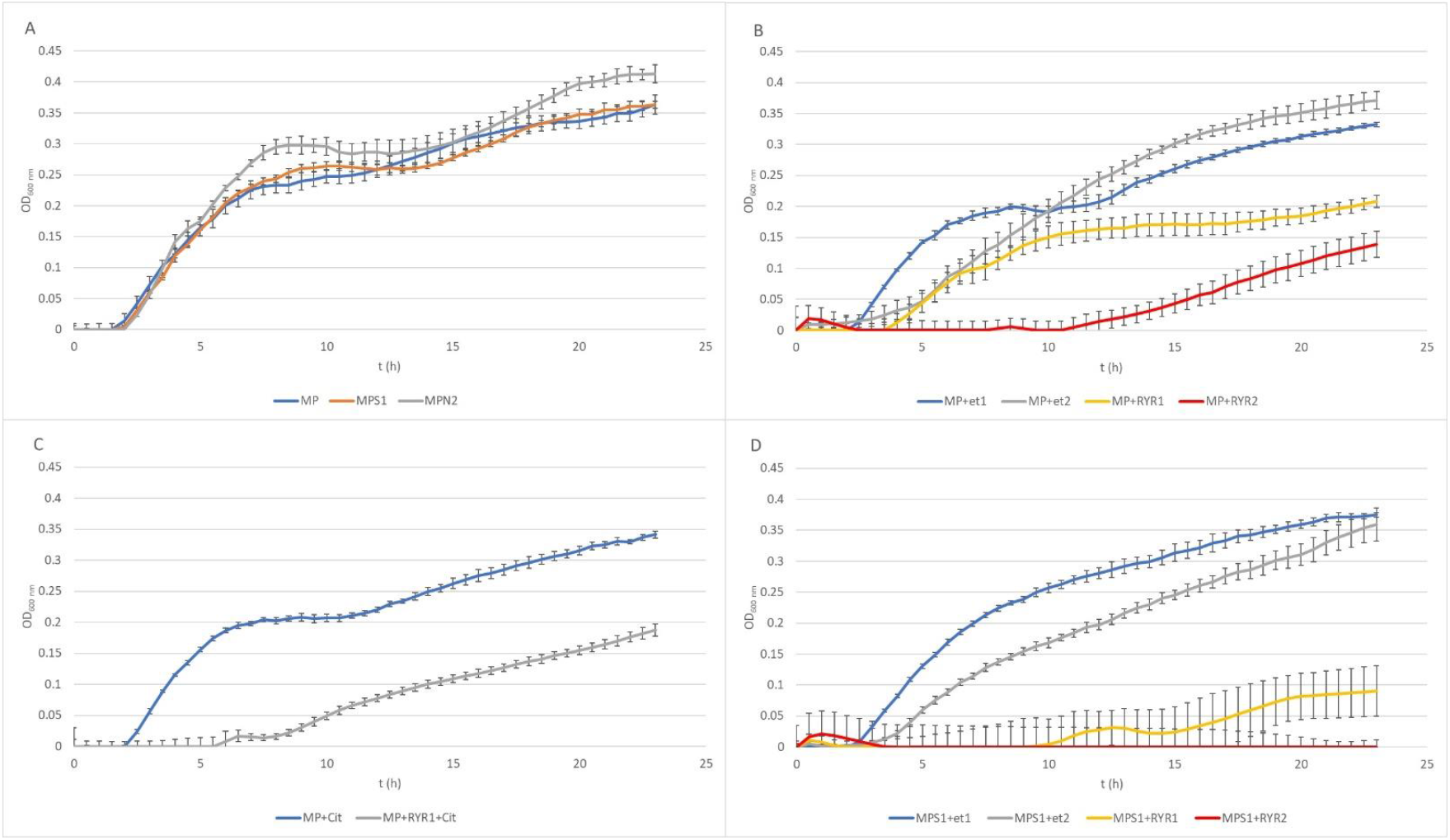
Growth curves of *B. subtilis* using different compositions of culture medium (for the experiment design see Table II) Graph A shows control experiment in MP medium and demonstrates the influence of NaCl to 1.3 % w/w (MPS1) and pickling nitrite salt to 2.0 % w/w (MPN2); Graph B shows the effect of ethanol at concentrations of 1.92 and 3.84 % v/v to MP medium (MP +Et1, MP + Et2) and addition of RYR ethanol extract to 1.92 and 3.84 % v/v (MP+ RYR1, MP+RYR2) Graph C shows the citrinin effect independently (MP+cit) or in combination with RYR extract (MP+RYR1+cit). Graph D shows the combined influence of NaCl (1.3 % w/w) and ethanol (1.92 and 3.84 % v/v); (MPS1+ Et1, MPS1 +Et2) or RYR ethanol extract (2 and 4 % v/v); (MPS1+ RYR1, MPS1 +RYR2).

### 3.3. RYR extract analysis

The absorption spectrum of the RYR extract is shown in Fig. 5. Values of absorption found by spectrophotometric analysis at 390, 470 and 500 nm, corresponding to assumed absorption maxima of yellow, orange and red pigments, were 98, 58 and 70, respectively. Monascin (yellow), rubropunctatin (orange) and rubropunctamine (red) were identified in the RYR extract by UHPLC analysis, (Fig. 6) while their analogs with seven carbon side chains i.e. ankaflavin (yellow), monascorubrin (orange) and monascorubramine (red) were not detected; neither was citrinin or monacolin K. The detected yellow pigments were quantified as monascin equivalents (1220 mg/L), orange pigments as rubropunctatin equivalents (336 mg/L) and red pigments as rubropunctamine equivalents (408 mg/L). However, other compounds labelled as yellow I and red I-VI were found and for their putative identification, HPLC-MS analysis and already published m/z data on different *Monascus* metabolites were used (for survey of the *Monascus* pigments data see Chen et al., 2019). While red I-red VI are probably rubropunctamine derivatives that were formed by the reaction of rubropunctatin with available amino group containing compounds, yellow I was identified as monascuspiloin (m/z 360.4).

**Fig. 5.**
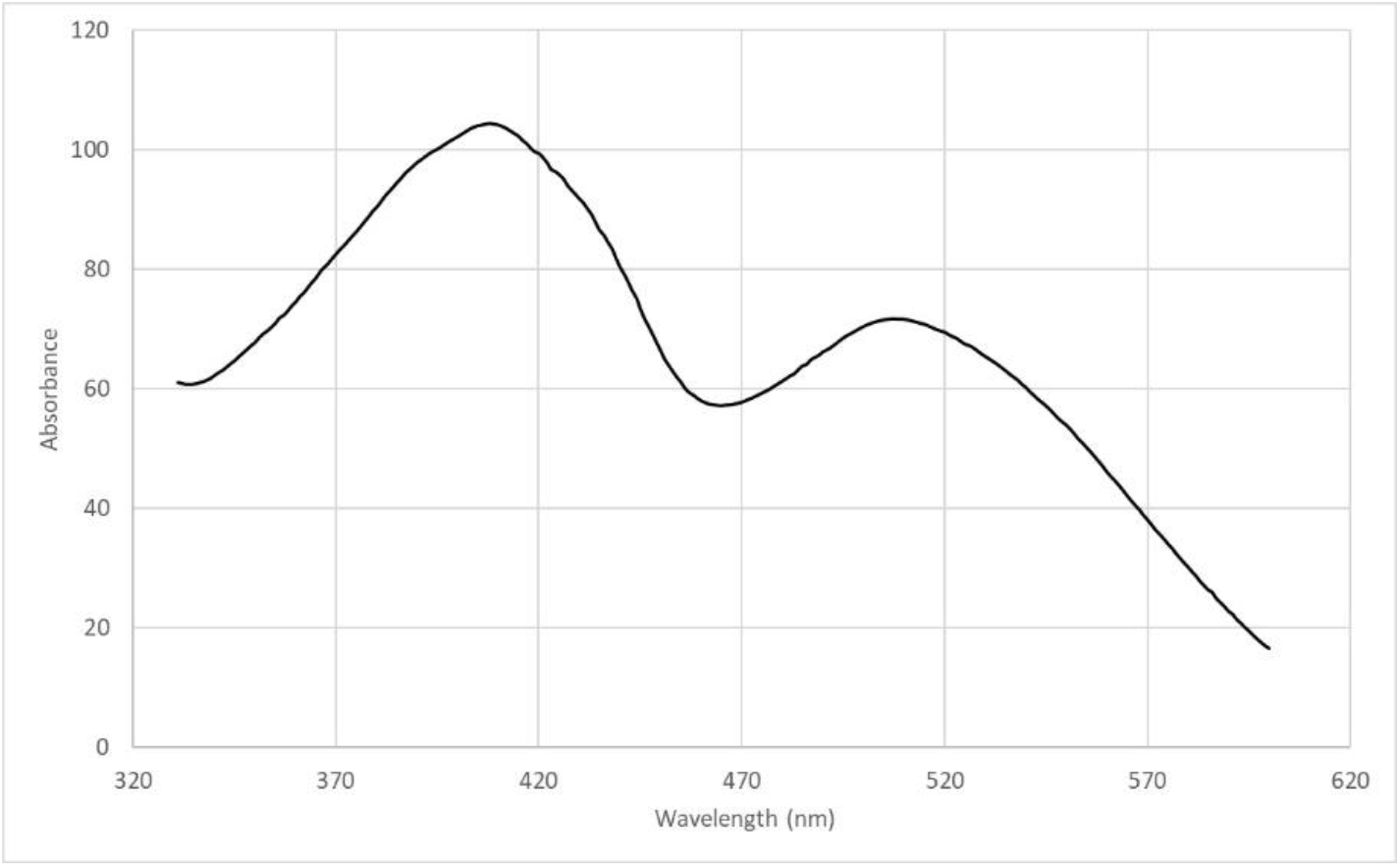
Absorption spectrum of the RYR extract.

**Fig. 6.**
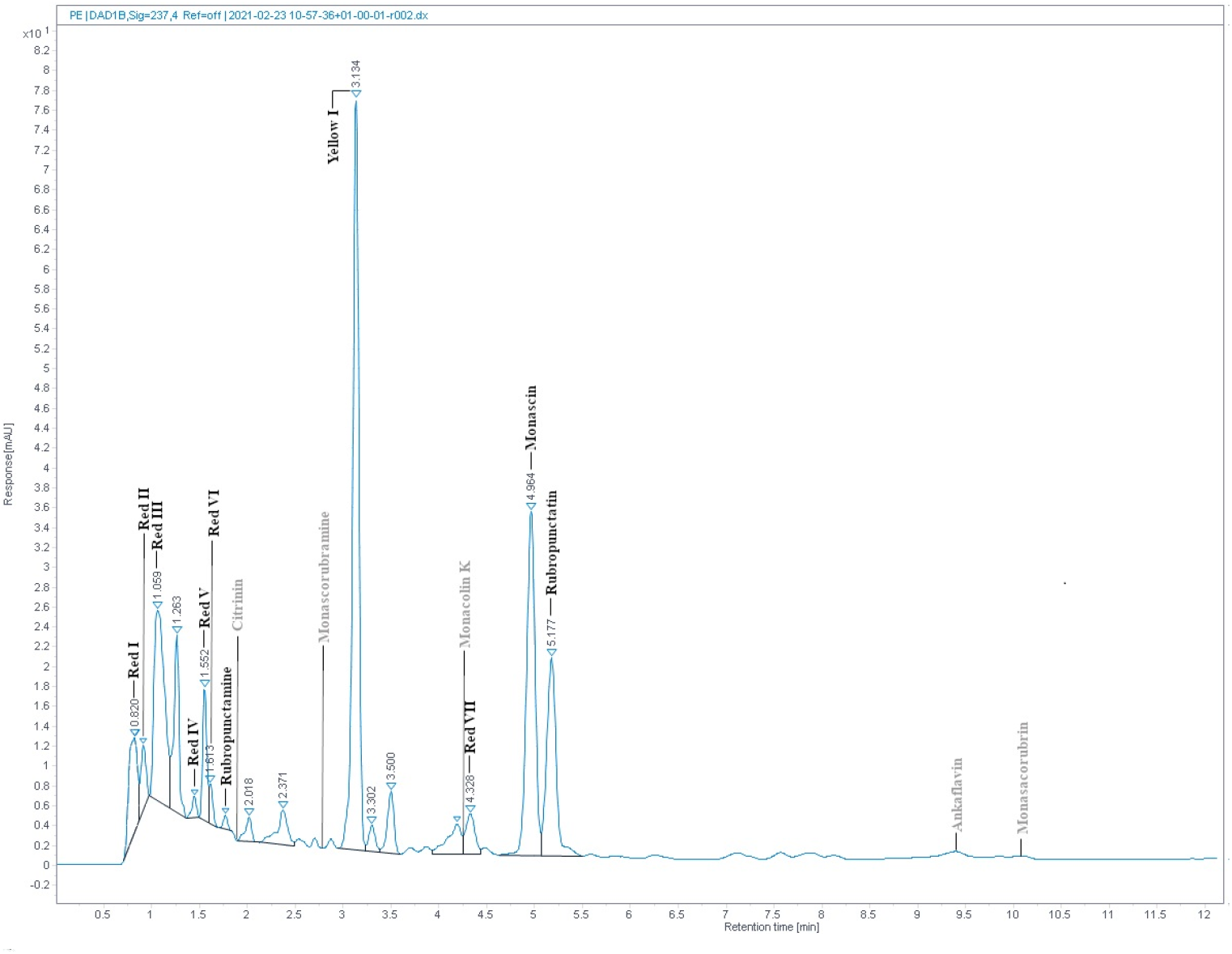
RYR extract chromatogram from UHPLC analysis. Detected compounds are given in black while expected but undetected compounds are given in grey colour. Conditions of UHPLC analysis are given in the section 2.3.

The RYR extract was added to TYA and MP medium to concentrations of 2 and 4 % v/v, respectively, and the concentrations of pigments added to the medium are shown in Table IV. The water extract of the RYR had a pH 4.9 but the pH of TYA or MP medium was 6.8 or 7.0. To determine whether the pigment profile was the same as the original RYR extract in the culture medium, samples of medium containing RYR mixed in a 1:1 volume ratio were analyzed by UHPLC. As expected, rubropunctatin was absent in the samples while monascin and monascuspiloin remained (data not shown).

**Table IV.** **Concentration of pigments quantified as monascin, rubropunctatin and rubropunctamine equivalents added to TYA and MP media** *Medium codes are the same as used in Tables I and II.

| Medium code* | Yellow pigments<br>(concentration equivalent to Monascin in mg/L) | Orange pigments<br>(concentration equivalent to Rubropunctatin in mg/L) | Red pigments<br>(concentration equivalent to Rubropunctamine in mg/L) |
| --- | --- | --- | --- |
| **TYA + RYR**<br>**TYAS1+ RYR** | 24.4 | 6.7 | 8.2 |
| **MP+ RYR1**<br>**MPS1 + RYR1** | 24.4 | 6.7 | 8.2 |
| **MP+ RYR2**<br>**MPS1+RYR2** | 48.8 | 13.4 | 16.4 |

## 4. Discussion

Nitrite and nitrate addition to meat products, together with NaCl, is traditional in European countries and is considered to be of low impact on human health even if cancerogenic nitrosamines can be formed in the acidic environment of the human stomach after ingestion of nitrite/nitrate containing food (Honikel, 2008; EFSA Panel, 2017). It is believed that the benefits of stable red colour, antioxidant and antimicrobial effects of nitrite/nitrate outweigh potential risks. Nevertheless, within this study it was found that addition of nitrite or nitrate alone, to the permitted concentration, did not suppress spore germination. Similar observations were documented in other studies for the germination of *Clostridium perfringens* (Labbe and Duncan, 1970), *Clostridium botulinum* (Sofos et al., 1979) or cheese associated clostridia including *C. beijerinckii* (Ávila et al., 2014) spores. However, combined effect of nitrite with NaCl tested after addition of the pickling salt both at the standard concentration of pickling salt (2 % w/w) and at the level recommended for low salt products (1.3 % w/w) was efficient in inhibiting the germination of *C. beijerinckii* spores but had no effect on germination of *B. subtilis* spores. However, total suppression of germination of *C. beijerinckii* spores was also achieved after addition of RYR extract to TYA medium, to a concentration of 2 % v/v while total suppression of germination of *B. subtilis* spores was only achieved after addition of 4 % v/v RYR extract to MP containing 1.3 % w/w NaCl. These results suggest that the RYR extract might substitute for nitrite salts in inhibiting germination of *Clostridium* spores.

Within the study, the ethanol effect on bacterial spore germination was confirmed (Setlow et al., 2002) as well as the effect of NaCl (Nagler et al., 2014) and the synergistic effect of different agents (Nerandzic et al., 2015). Even if the RYR extract did not contain citrinin, its effect at 2000 µg/L (the permitted concentration of citrinin in RYR food supplements in the EU until 2019) was tested, but if applied independently, had no effect on spore germination.

*Monascus* sp. DBM 4361, used for the preparation of RYR, was not classified on the species level but it might be *M. pilosus* because its characteristics, i.e., no citrinin production and formation of both conidia and ascospores, corresponds with an already described strain, *M. pilosus* MS-1, also isolated from the red yeast rice (Feng et al., 2016). The absence of citrinin production was also found in *M. pilosus* NBRC4520 (Higa et al., 2020). In the RYR extract, there were found three of six iconic *Monascus* pigments; in particular monascin, rubropunctatin and rubropunctamine, together with rubropunctamine complexes with different amino group-containing compounds and monascuspiloin, a yellow pigment with a structure similar to monascin that has already been described as the metabolite of *M. pilosus* M93 (Chen et al., 2012). Interestingly, only monascin and rubropunctatin, i.e. pigments with a shorter five carbon side chain, but not their analogs (ankaflavin and monascorubrin with seven carbon side chains) were found. During the biosynthesis of pigments by *M. ruber* M7 (Chen et al., 2017), MrPigJ and MrPigK subunits of fatty acid synthase were found as the proteins responsible for the integration of β-ketooctanoic or β-ketodecanoic acid moieties into the structure of the pigments. However, the selection of the particular fatty acid moiety was considered to be random or not yet understood. Homologous genes to *MrPigJ, MrPigK* were described in *M. purpureus* (*MpFasA2, MpFasB2*) (Balakrishnan et al., 2013) and other *Monascus* strains (Guo et al., 2019) but not in *M. pilosus*. Even in the newest review of azaphilone biosynthesis (Pavesi et al., 2021), factors that determine the selection of particular fatty acid moieties are not described.

For the detection of substances in *Monascu*s extracts, it is not sufficient to use the absorption spectrum or to determine absorbance values of the extract at the absorption maxima of individual pigments, typically at 390, 470 and 500 nm. It is really necessary to analyze extracts by HPLC or other analytical method (cf. Fig.5 and Fig.6). Notwithstanding, the determination of individual compounds in *Monascu*s extracts is difficult because it depends inter alia on the pH of the extract (Shi et al., 2016).

After the addition of the RYR extract to medium, the original pH of the RYR extract changed from 4.9 to 6.8 or 7.0, resulting in the reaction of rubropunctatin with available amino group containing compounds, such as amino acids. It is possible that this reaction might contribute to the suppression of spore germination in the medium. Amino acids such as L-alanine or glycine are known germinants (factors stimulating germination) of bacterial spores (Setlow, 2014; Bhattacharjee et al., 2016) and if their amount was decreased it might affect germination. In addition, it was reported that *Monascus* red pigment derivatives had pronounced effects on the growth of Gram positive bacteria, including *B. subtilis* (Kim et al., 2006). The assumed cause of inhibition was adsorption of pigment derivatives onto the surface of cells, limiting oxygen uptake, where their MIC values were found to be 4-8 µg/L. In our assay, rubropunctatin and a mixture of red pigment derivatives, to concentrations of 13.4 mg/L and 16.4 mg/L, respectively, were added to MP culture medium used for outgrowth of *B. subtilis* spores (see Table IV) and *B*.*subtilis* was cultured under aerobic conditions; the above aerobic effect (Kim et al., 2006) might also apply here. Orange and red *Monascu*s pigments, at concentrations of 10-20 mg/L, inhibited growth of Gram negative bacteria (Vendruscolo et al., 2014), which corresponds with our findings. Yellow pigments monascin and monascuspiloin, detected in the RYR,, were found to have anticancerogenic effects (Akihisa et al., 2005; Chen et al., 2012; Chiu et al., 2012) but their antibacterial effect was never tested.

## Acknowledgement

Financial support from specific university research (MEYS No 8-SVV/2021).

